# Temporal and spatial variation in distribution of fish environmental DNA in England’s largest lake

**DOI:** 10.1101/376400

**Authors:** Lori Lawson Handley, Daniel S. Read, Ian J. Winfield, Helen Kimbell, Harriet Johnson, Jianlong Li, Christoph Hahn, Rosetta Blackman, Rose Wilcox, Rob Donnelly, Amir Szitenberg, Bernd Hänfling

## Abstract

Environmental DNA offers great potential as a biodiversity monitoring tool. Previous work has demonstrated that eDNA metabarcoding provides reliable information for lake fish monitoring, but important questions remain about temporal and spatial repeatability, which is critical for understanding the ecology of eDNA and developing effective sampling strategies. Here, we carried out comprehensive spatial sampling of England’s largest lake, Windermere, during summer and winter to 1) examine repeatability of the method, 2) compare eDNA results with contemporary gill-net survey data, 3) test the hypothesis of greater spatial structure of eDNA in summer compared to winter due to differences in water mixing between seasons, and 4) compare the effectiveness of shore and offshore sampling for species detection. We find broad consistency between results from three sampling events in terms of species detection and abundance, with eDNA detecting more species than established methods and being significantly correlated to rank abundance determined by long-term data. As predicted, spatial structure was much greater in the summer, reflecting less mixing of eDNA than in the winter. For example Arctic charr, a deep-water species, was only detected in deep, mid-lake samples in the summer, while littoral or benthic species such as minnow and stickleback were more frequently detected in shore samples. By contrast in winter, the eDNA of these species was more uniformly distributed. This has important implications for design of sampling campaigns, for example, deep-water species could be missed and littoral/benthic species overrepresented by focusing exclusively on shoreline samples collected in the summer.

## Introduction

Rapid monitoring of biodiversity for conservation, management or assessing the impact of anthropogenic pressures is frequently difficult to achieve using established methods. This is particularly relevant for fish in lake ecosystems, as no established method is suitable across all lake sizes and depths: electrofishing is unsuitable for large, deep lakes; gill-netting under-records species restricted to very shallow water and is destructive; and hydroacoustics has low efficacy in shallow lakes and is unable to identify species. Environmental DNA (‘eDNA’), which is released by organisms into their environment in the form of shed cells, excretia, gametes or decaying matter (Taberlet, Coissac, Hajibabaei, & Rieseberg, 2012), is promising as a complementary or alternative method for monitoring fish in lakes (Jerde, Mahon, Chadderton, & Lodge, 2011; Civade et al., 2016; Hänfling et al., 2016; Lacoursière-Roussel, Côté, Leclerc, & Bernatchez, 2016; Valentini et al., 2016; Evans & Lamberti, 2017; Evans et al., 2017; Hering et al., 2018;) and PCR-based metabarcoding of eDNA has tremendous potential for monitoring entire ecological communities (see, for example, Bohmann et al., 2014; Lawson Handley, 2015; Valentini et al., 2015; Deiner et al., 2017 for reviews).

Although eDNA metabarcoding is still in its infancy, a great deal of progress has been made very recently and a number of studies have demonstrated that it can effectively describe fish communities in lentic (Civade et al., 2016; Hänfling et al., 2016; Evans et al., 2017; Valentini et al., 2016; Klymus, Marshall, & Stepien, 2017), lotic (Civade et al., 2016; Shaw et al., 2016; Valentini et al., 2016) and marine environments (e.g. Thomsen et al., 2012, 2016; Miya et al., 2015; Port et al., 2016; Andruszkiewicz et al., 2017; O’Donnell et al., 2017; Yamamoto et al., 2017). eDNA metabarcoding consistently outperforms established methods for detection of fish species (e.g. Thomsen et al., 2012, 2016; Miya et al., 2015; Civade et al., 2016; Hänfling et al., 2016; Port et al., 2016; Shaw et al., 2016; Valentini et al., 2016; Andruszkiewicz et al., 2017; Yamamoto et al., 2017) and is at least semi-quantitative, correlating with data from established surveys and providing estimates of (at least relative) abundance (Evans et al., 2015; Hänfling et al., 2016; Port et al., 2016; Andruszkiewicz et al., 2017; O’Donnell et al., 2017).

In our previous work we demonstrated that eDNA metabarcoding has huge potential for describing fish community structure in lakes (Hänfling et al., 2016). Water samples were collected in January 2015 from Windermere, the largest lake in England, assayed by eDNA metabarcoding of mitochondrial 12S and cytochrome b (CytB) and compared to data from gill-netting and hydroacoustic surveys. Windermere is arguably the most intensively studied lake in the UK, with data on fish populations, physicochemical and other biological properties collected over many years and regular monitoring of fish populations since the 1940s (Maberly et al., 2011; Winfield, Fletcher, & James, 2016). Firstly, 14 of the 16 species ever recorded in Windermere were detected using eDNA compared to only four species detected in an extensive gill-net survey carried out four months prior to eDNA sampling. Interestingly, more species were detected in shallower water, and 12 of the 16 species were detected in just 6 spatially-close shoreline samples. This suggests that eDNA could accumulate at the shoreline and that shoreline sampling could be adequate for detection of most species but more rigorous sampling along the shoreline is needed to investigate this further. Secondly, depth transects revealed that most species’ eDNA was distributed throughout the water column, but eDNA of the deep-water species Arctic charr (*Salvelinus alpinus*) was only detected at the deepest sampling points, indicating that surface water sampling may be ineffective for some species in deep lakes. Thirdly, we found a strong spatial signal in the distribution of eDNA from species that prefer the more mesotrophic conditions of the lake’s North Basin, compared to those that are associated with the more eutrophic conditions of the South Basin. This indicates that eDNA provides a contemporary signal, at least to some extent, of the fish distribution, and that eDNA is promising for ecological assessment of water bodies. Moreover, eDNA abundance data consistently correlated with rank abundance estimates from established surveys, demonstrating, together with other studies (e.g. Evans et al., 2015; Thomsen et al., 2016) that at least semi-quantitative estimates could potentially be obtained from eDNA data. Critical questions remain about the spatial and temporal distribution of eDNA in order to better understand the ecology of eDNA and design the most effective strategy for future monitoring programmes. For example, 1) how does eDNA distribution vary between seasons, 2) is shoreline sampling more effective than offshore sampling for species detection, and 3) how do abundance estimates from eDNA compare to those from established methods carried out at the same time? We explore each of these questions in the current study, by adding data from summer and winter sampling campaigns on Windermere.

There are several reasons why eDNA distribution might vary at different times of the year, including patterns of water mixing, fish behaviour and distribution, and different rates of DNA degradation. In our previous study, water samples were collected in winter, when lakes are unstratified and water is extensively mixed in the vertical dimension (Hänfling et al., 2016). During summer, deeper lakes are stratified and show strong vertical gradients in temperature, while most fish species are also likely to be present and active. Assessing temporal variability is crucial for determining the repeatability of eDNA based methods, but seasonality of eDNA signal has so far been little explored (but see e.g. Sigsgaard et al., 2017; Tillotson et al., 2018). Here, we test the hypothesis that there will be a stronger spatial structure of eDNA in the summer compared to winter.

Shoreline sampling is an attractive option for biodiversity monitoring as it avoids the costs, specialist training, access to equipment, and health and safety considerations associated with boat-based work. To investigate whether shoreline sampling is adequate for detection of most species, we collected samples from the entire perimeter of Windermere and compared shoreline samples to those from offshore transects. We hypothesized that more species will be detected in the shoreline samples, and that fewer samples will be needed for species detection, relative to offshore samples. We also predict this effect will be greatest in the summer, due to greater spatial structure as discussed above.

Obtaining accurate estimates of species abundance and biomass remains arguably the greatest challenge for eDNA applications due to the large number of factors that influence DNA dynamics (Barnes & Turner, 2015; Barnes et al., 2014) and the many opportunities for bias during sampling, lab and bioinformatics workflows (Valentini et al., 2016). In our previous study we tested the efficacy of both sequence read count and site occupancy (i.e the proportion of samples in which a species was detected) for assessing relative abundance (Hänfling et al., 2016). Encouragingly, both measures were significantly correlated to rank abundance, but comparative established survey data was based on historical datasets (up to September 2014) and expert opinion, and further work is needed to determine how robust eDNA is for estimating abundance. To explore this further, and ensure that comparisons between methodologies are as robust as possible, we performed eDNA sampling at the same time as the annual gill-net survey in September 2015.

In the present study, eDNA samples were collected from Windermere along three offshore transects and the entire shoreline in September 2015 and January 2016, and along ten depth profiles (September only), then data combined with that from January 2015 (Hänfling et al., 2016) in order to: 1) examine the temporal repeatability of eDNA metabarcoding for lake fish communities across seasons (summer and winter) and years (2015-2016), 2) compare eDNA results with data from gill-net and hydroacoustic surveys carried out at the same time of sampling, 3) test the hypothesis of greater spatial structure of eDNA in summer compared to winter due to water stratification in summer and breakdown in winter, and 4) robustly compare the effectiveness of shore and offshore sampling locations for species detection.

## Materials and Methods

### Study site

Windermere is 16.9 km in length, with a surface area of 1480 ha. The lake is divided into two separate basins: North and South Basin, by a shallow area with islands. North Basin is classed as mesotrophic and has a maximum depth of 64 m. South Basin is more eutrophic and has a maximum depth of 44 m. Lake stratification typically begins in April and persists to November, during which period, the thermocline usually occurs at a depth of between 10 and 20 m.

### Established surveys

Gill-netting surveys were carried out between 1st-3rd September 2015 at five sites (including a surface site directly above a deep-water bottom site) in each of the two Windermere basins, as described in detail by (Winfield et al., 2016). We previously summarised fish species presence and abundance for Windermere based on a literature review and IJW’s expert opinion (Hänfling et al., 2016). Each of the 16 previously recorded species was assigned a relative long-term abundance score ranking from 1 (most common) to 16 (least common, Table S1). The same rank classification is adopted in the present study.

### eDNA Sampling

Two sampling events were carried out in Windermere during summer (9th-13th September 2015) and winter (26th-28th January 2016). Two-litre water samples, comprised of 5 × 400 ml pooled subsamples, were collected as described in (Hänfling et al., 2016). Offshore samples were collected from a boat using a Friedinger sampler, along three transects with approximately 1 km sampling interval between sites, at the same locations sampled in (Hänfling et al., 2016) (Figure 1, Table S2). Transect 1 follows the 5 m depth contour (green dots in Figure 1, N = 16), transect 2 follows the 20 m depth contour, (red circles, N = 14), and transect 3 follows the lake midline (blue and purple circles, N = 15). This sampling scheme covered 7 of the 10 sites that are used for annual gill-net surveys and samples were also collected at the remaining three gill-net sites (black triangles, Figure 1). Water samples were collected at approximately half the water depth (i.e. nominally in the metalimnion) at each of the offshore sites. During the summer sampling campaign, samples were collected at 10 sites (purple circles, Figure 1) from the midline transect from the surface (epilimnion) and approximately 2 m above the lake bottom (hypolimnion) in order to investigate the effects of stratification on eDNA distribution (N = 21). The Friedinger sampler was sterilised between samples by washing in a 10% commercial bleach solution (containing ~3% sodium hypochlorite) followed by 10% microsol detergent (Anachem, UK) and rinsed with purified water. Sampling blanks were collected after approximately every 8 samples by running 2 L of purified water through the Friedinger sampler after sterilisation (N = 9 for September 2015 and 7 for January 2016). Shore samples were collected directly into sterile 2 L plastic bottles. The 40 shoreline sample sites were approximately 1 km from each other and aligned with the offshore transects as far as this was possible based on accessibility. All samples were stored on ice in a cooler prior to filtration. The total number of samples excluding blanks was 108 (N = 69 offshore and 40 shore) in September 2015 and 87 (N = 47 offshore and 40 shore) in January 2016 in addition to the 78 (72 offshore and 6 shore) samples collected by (Hänfling et al., 2016).

**Figure 1.**
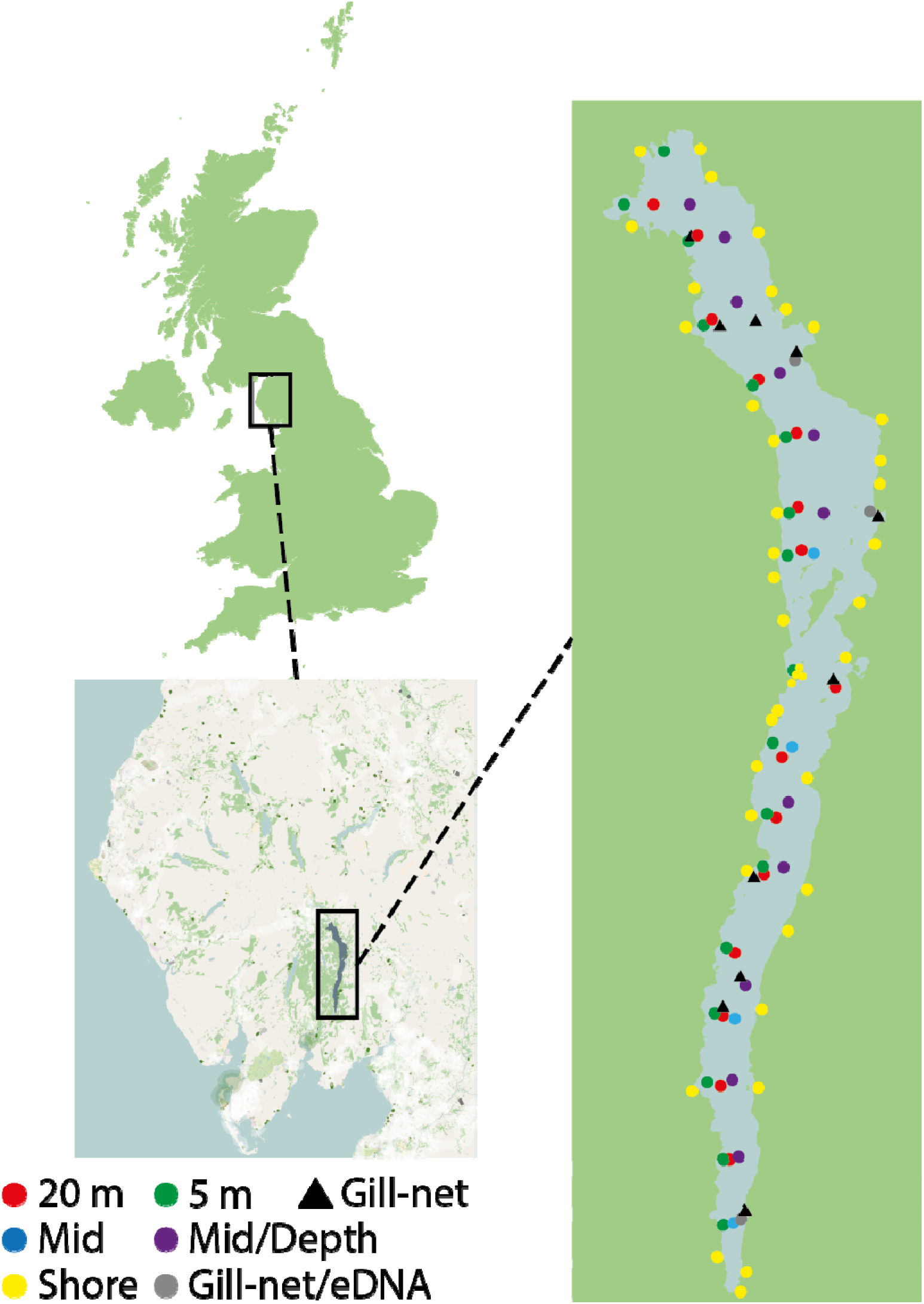
Distribution of shore and offshore sampling sites in Windermere. Coloured dots correspond to the following sample types: red, 20 m offshore transect; green, 5 m offshore transect; orange, gill-net survey sites; blue, midline offshore transect sites; purple, sites on midline transect where depth profiles were taken in September 2015; yellow, shore line sites; black triangles, additional sites adjacent to gill-net survey sites that were separate from the main transects. Sample coordinates are provided in Table S2.

### DNA capture, extraction and library preparation

Water filtration was carried out at the Freshwater Biological Association laboratories at Windermere, within 8 hours of collection, in a laboratory that does not handle fish. Samples were filtered through 0.45 μm cellulose nitrate filters and pads (47 mm diameter, Whatman, GE Healthcare, UK) using Nalgene filtration units in combination with a vacuum pump. In a previous study we demonstrated that 0.45 μm cellulose nitrate filters are suitable for fish metabarcoding, with low variation and high repeatability between filtration replicates (Li, Lawson Handley, Read, & Hänfling, 2018). All filtration equipment was sterilized in 10% commercial bleach solution for 10 minutes, followed by rinsing in 10% microsol and purified water after each filtration. Filtration blanks were run before the first filtration and then after every ten samples (N = 35), in order to test for possible contamination at the filtration stage. Filters were stored in sterile 50 mm petri dishes, sealed with parafilm, at −20°C until DNA extraction. DNA was extracted from filters using the PowerWater DNA Isolation Kit (MoBio Laboratories, Carlsbad, USA) and the manufacturer’s protocol.

DNA samples were amplified at two mitochondrial regions: 12S rRNA (12S, 106 bp, Kelly, Port, Yamahara, & Crowder, 2014; Riaz et al., 2011) and cytochrome b (CytB, 414 bp, Kocher et al., 1989) using 16 individually-tagged forward primers and 24 individually-tagged reverse primers, with one-step library preparation as described in (Hänfling et al., 2016) but with minor modifications. PCR reactions contained 0.5 μM each primer, 200 mM dNTPs, 12.5 μL Q5^®^ High-Fidelity 2X Master Mix (New England Biolabs) and <1000 ng template DNA. PCR profiles were as follows: 98°C for 30 seconds followed by 35 cycles of 98 °C for 10 seconds, 58 °C (12S)/50°C (CytB) for 15 seconds and 72 °C for 20 seconds, and a final extension step of 72°C for 5 minutes. PCR negative controls included each primer at least once (N = 40). Each PCR reaction was carried out in triplicate and pooled in order to reduce potential bias through stochastic variation during the PCR step. PCR products were checked on ethidium bromide-stained agarose gels. Each set of samples was normalised for concentration across the samples using the Life Technologies SequalPrep Normalization Plate Kit and subsequently pooled to make a single sequencing library for each assay (12S and CytB). Samples were split across two libraries per locus (hence 4 libraries in total). Each library was quantified using the Qubit HS DNA Quantification Kit (ThermoFisher) and sequenced on an Illumina MiSeq using V3 2 × 300 bp chemistry at 8 pM concentration including a 10% addition of PhiX.

### Bioinformatics and data analysis

Raw read data for all four libraries have been submitted to NCBI (BioProject: <u>PRJNA482277</u>, SRA Study:<u>SRP154799</u>. https://www.ncbi.nl?n.nih.gov/Traces/studv/?acc=SRP154799). Bioinformatics was carried out using a custom-made reproducible metabarcoding pipeline (metaBEAT v0.97.9) with a custom reference database of 67 European freshwater fish species as described in our previous studies (Hänfling et al., 2016; Li et al., 2018). A brief overview of steps taken in the bioinformatics pipeline is provided in the Supplementary Material. To assure full reproducibility of our bioinformatics steps, the reference databases and Jupyter notebooks for data processing have been deposited in a dedicated GitHub repository (https://github.com/HullUni-bioinformatics/Handley_et_al_2018).

Filtered data were summarised in two ways for downstream analyses: the number of sequence reads per species divided by the total number of reads per sample (normalised read counts) and the proportion of sites occupied by a species (site occupancy). To reduce the possibility of false positives, we only regarded a species as present at a given site if its sequence frequency exceeded a certain threshold level (proportion of all sequence reads in the sample) which was established in (Hänfling et al., 2016) as 0.1% and 0.2% for 12S and CytB respectively.

The relationship between eDNA data and data from established surveys (rank abundance by numbers or rank biomass based on long term expert opinion, and biomass estimates from September 2015 gill-net surveys) was investigated by calculating Spearman’s Rho (for rank correlations) and Pearson’s Product-moment correlation coefficient (for biomass) in R v3.1.3 (R Core Team 2017). Data was plotted by fitting a smoothed linear model with the function geom_smooth(model=lm) in ggplot2 (Wickham, 2009).

The direct comparison between eDNA data and contemporary gill-net data was based on 10 sites that had complete data for both eDNA and gill-netting surveys. Only species detected in the gill-netting surveys were included in this analysis. Normalised read counts per species were calculated by summing the total read count per species for the 10 sites, and dividing by the total read count for these species and sites. Similarly, biomass estimates from gill-netting data were normalised by dividing the total biomass for a species by the total biomass for all species.

The analyses were repeated for both loci and on different hierarchical levels: i) all Windermere samples, ii) basins (North and South), iii) transects within basins, and iv) depth profiles within transect to investigate the spatial and temporal variation in eDNA distribution. Finally, sample-based rarefaction (Gotelli & Colwell, 2010)was used to determine the number of samples needed to accurately represent the species assemblage. Rarefaction was performed using the functions *rich* and *rarc* with 499 randomizations in the R package Vegan v2.4-4 (Oksanen, 2015) for both loci, shore and offshore samples, and summer and winter.

## Results

### Established surveys

Gill-netting surveys detected five species and a single hybrid individual (roach, *R. rutilus x* common bream, *A. brama*, Table S1). Similar numbers of fish were caught in the North and South Basins (N = 681 and 709 respectively). Perch, *Perca fluviatilis*, was by far the most abundant species in both basins (N = 517 and 644 in North and South Basins respectively) in terms of both numbers and biomass (Table S1). Roach was also found in both basins but at higher numbers in the North Basin (N = 161 compared to N = 61 for the South Basin). Atlantic salmon, *Salmo trutta* (N = 2) and pike, *Esox lucius* (N = 1) were found in the North Basin only, while common bream (N = 2) was detected at a single site in the South Basin.

### Library quality, raw data and controls

Libraries generated between 1.175 and 2.84 Gbp data and had average %≥Q30 scores of 75.40 to 75.54 (CytB 2.84 Gbp, Q30 75.40, 12S, 1.75 Gbp, Q30 75.54). Sequencing libraries contained on average 18.37 million raw reads (September: 17.81 million for CytB, 30.25 million for 12S; January 5.54 million for CytB, 20.78 million for 12S), of which, an average of 13.46 million reads passed filter (September: 16.62 million for CytB, 19.77 million for for 12S; January: 4.62 million for CytB, 11.64 million for 12S). After quality filtering and removal of chimeric sequences, the average read count per sample (excluding controls and samples sequenced for other projects) over all four libraries was 38,124 (average total read counts by library: September: 85,0003 for CytB, 32,797 for 12S; January: 22,223 for CytB, 12,474 for 12S). The average number of fish sequences per sample over all four libraries was 30,012 (average fish read counts per library: September: 84,560 for CytB, 20,254 for 12S; January: 10,745 for CytB, 4,488 for 12S). A similar average January fish read count was obtained in our previous dataset from January 2015 (8,219 for CytB, 6,842 for 12S, Hänfling et al., 2016) indicating lower fish read count in the winter months. Full run metrics are provided in Table S3.

Negligible amounts of contamination were found in the January samples for both loci, with a total of just 19 reads over 30 negative control samples for 12S and 7 reads over 32 samples for CytB. Contamination can therefore be confidently ruled out for these samples. In the September CytB library, 324 reads were detected over 14 PCR negatives and 195 reads were detected in 22 sample and filtration blanks. These reads were almost exclusively assigned to perch, and the maximum number of reads per sample was 47. This indicates a very low level of perch contamination in the September CytB dataset. In the September 12S library, a total of 107 reads was detected over 35 PCR negative controls. Roach was detected in 7, perch detected in 6 and minnow, *Phoxinus phoxinus*, detected in 5 PCR negatives, but the maximum number of reads per sample, per species was just 12, suggesting contamination is negligible at the PCR stage. Nine of 23 sample/filtration blanks in this library had zero sequence reads, however notable evidence of contamination (i.e. sequence reads in the order of 1000) was found in the remaining 14 blanks. A total of nine species was detected, three of which (tench, *Tinca tinca*; roach and brown trout, *Salmo trutta*) are known to be present in Windermere. It is therefore important to bear in mind that the read counts for these species may be inflated in the actual samples for this library. The other six species detected (Crucian carp, *Carassius carassius*; gudgeon *Gobio gobio*, common bleak, *Alburnus alburnus*, mudminnow, *Umbra pygmaeus*; common carp, *Cyprinus carpio* and chub, *Squalius cephalus)* have not been recorded by established surveys in Windermere. However common carp and mudminnow were detected with 12S at one site each in our January 2015 sampling (Hänfling et al., 2016). Because of the ambiguity introduced by this contamination issue, we restrict the results to species that have been previously confirmed in the lake.

### Species detection with eDNA

Detection of previously-recorded species was generally comparable between loci and seasons, with some exceptions. Fourteen of the 16 species recorded with established methods (i.e. all species excluding the lampreys: river lamprey, *Lampetra fluviatilis* and sea lamprey, *Petromyzon marinus*) were detected in total (Figure 2, S1 and S2). All of these species were detected in both winter sampling campaigns with 12S. Eight species (perch, roach, Arctic charr, pike, brown trout, eel, *Anguilla anguilla*; bullhead, *Cottus gobio*; and common bream) were detected with both markers in both basins in all three sampling events (Figure 2, S1 and S2). All five species detected in the September 2015 gillnet survey were detected in the eDNA data.

**Figure 2.**
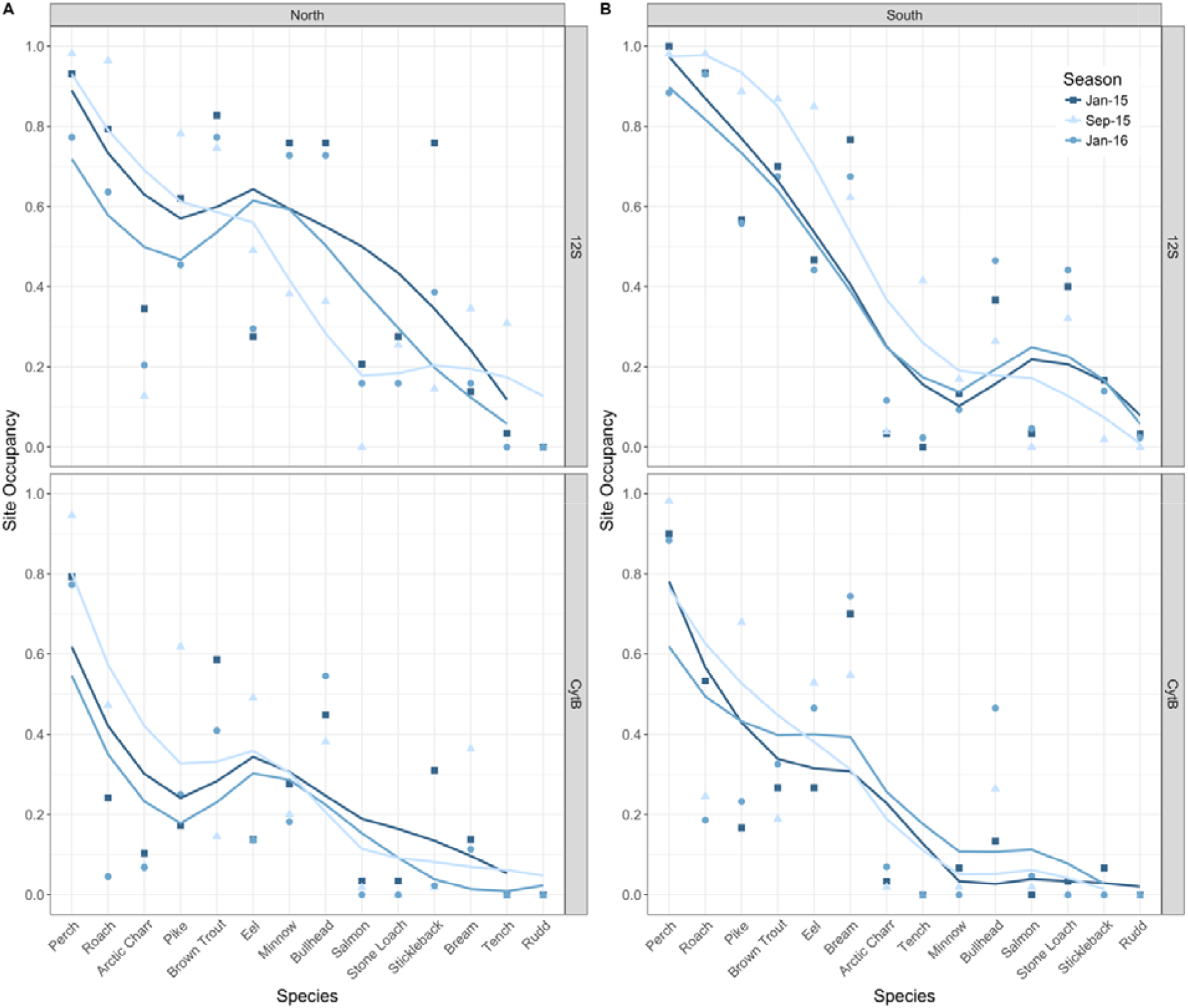
Species detection in January 2015, September 2015 and January 2016 based on site occupancy, for Windermere North (A) and South (B) Basins. Species are ordered according to their long term rank within the basins, with perch the most abundant and rudd the least abundant in both basins. Smoothed curves were fitted with a linear model. See Table 1 for results of correlations.

Some differences were observed between markers (Figure 2, S1 and S2). Tench and rudd, *Scardinius erythropthalmus*, were not detected with CytB. Occupancy for some species (e.g. roach) was consistently higher with 12S (Figure 2a and b, S1 and S2, a and b) than with CytB (Figure 2c and d, Figure S1 and S2, c and d). Stone loach, *Barbatula barbatula*, was detected in all three seasons and both basins with 12S, but was only detected in January 2015 with CytB. In general there was more consistency in site occupancy between sampling events with 12S than CytB. Note that the occupancy of tench in the September 2015 12S data could be inflated by contamination and should therefore be interpreted with caution.

There were also some notable differences in detection between seasons and between basins (Figure 2, S1 and S2). For example, detection of some species (e.g. pike, eel) was consistently greater in summer than in winter, while bullhead had higher detection rates in the winter (most notably in North Basin). Rudd, the rarest of the 14 detected species, was only detected in the winter (with 12S in the South Basin). Differences between basins, observed in previous work, were confirmed. In particular, common bream had higher site occupancy in the South Basin compared to North Basin in all seasons (Hänfling et al., 2016), but was more common in the North Basin and less common in the South Basin in summer compared to winter. Detection of Atlantic salmon was also higher in winter than in summer and in North Basin compared to South Basin.

### Correlations between eDNA and data from established surveys

In spite of the observed differences between loci and seasons discussed above, correlations between eDNA data and long term rank were highly consistent between seasons and between loci (Figure 2 and Table 1). Twenty-three of 24 correlations between eDNA data and long term rank were significant (Table 1). Similar results were found for both site occupancy and read count (Table 1). Spearman’s rho and corresponding *P* values were consistently higher for 12S than for CytB (12S rho = −0.695 to −0.905, *P* < 0.005; CytB rho = −0.584 to −0.795, *P* < 0.05).

**Table 1.** Results of Spearman’s Rank Correlations between Long Term Rank and Read Count (Proportion of total read count) and Site Occupancy.

| Sampling event | Locus | Basin | Read Count | Site Occupancy |
| --- | --- | --- | --- | --- |
| Jan 15 | 12S | North | $\rho = -0.710, S = 778, P = 0.006$ | $\rho = -0.758, S = 799.9, P = 0.002$ |
| Sep 15 | 12S | North | $\rho = -0.660, S = 755.33, P = 0.010$ | $\rho = -0.695, S = 771.35, P = 0.006$ |
| Jan 16 | 12S | North | $\rho = -0.612, S = 733.31, P = 0.020$ | $\rho = -0.733, S = 788.58, P = 0.003$ |
| Jan 15 | 12S | South | $\rho = -0.793, S = 816, P = 0.001$ | $\rho = -0.722, S = 783.45, P = 0.004$ |
| Sep 15 | 12S | South | $\rho = -0.818, S = 827.41, P = 0.0003$ | $\rho = -0.905, S = 866.91, P = 8.474e-06$ |
| Jan 16 | 12S | South | $\rho = -0.798, S = 818, P = 0.001$ | $\rho = -0.745, S = 794.12, P = 0.002$ |
| Jan 15 | CytB | North | $\rho = -0.422, S = 647.21, P = 0.132$ | $\rho = -0.584, S = 720.88, P = 0.028$ |
| Sep 15 | CytB | North | $\rho = -0.589, S = 723.18, P = 0.027$ | $\rho = -0.736, S = 789.84, P = 0.003$ |
| Jan 16 | CytB | North | $\rho = -0.536, S = 698.69, P = 0.048$ | $\rho = -0.633, S = 743.18, P = 0.015$ |
| Jan 15 | CytB | South | $\rho = -0.748, S = 795.5, P = 0.002$ | $\rho = -0.777, S = 808.73, P = 0.001$ |
| Sep 15 | CytB | South | $\rho = -0.747, S = 794.75, P = 0.002$ | $\rho = -0.795, S = 816.61, P = 0.0007$ |
| Jan 16 | CytB | South | $\rho = -0.681, S = 764.89, P = 0.007$ | $\rho = -0.707, S = 776.51, P = 0.005$ |

Sequence read counts were positively correlated to biomass of the five species detected in the September 2015 gillnet surveys for both 12S (Pearson’s product moment correlation coefficient *r* = 0.911 t = 3.837, df = 3, *P* = 0.031, Figure S3a) and CytB (*r* = 0.935 t = 4.572, df = 3, *P* = 0.019, Figure S3b).

### Spatial distribution of eDNA

We noted above that differences in spatial distribution were observed between North and South Basins for species such as common bream. Here we focus on the comparison of shore, offshore and depth transects along the entire lake (Figure 1) for summer (September 2015) and winter (January 2016). For perch, roach, pike, brown trout, eel, and tench, the distribution of eDNA is uniform between transects, and this observation is repeatable between seasons (Figure 3). By contrast strong spatial structuring was observed in the summer for some species. Most notably, Arctic charr was only detected in the offshore transects in the summer, and occupancy increased from the 5 m to midline transect (i.e. with depth), whereas in winter, this species was detected in all four transects (Figure 3). The reverse summer pattern was observed for minnow, bullhead, stone loach and three-spined stickleback, *Gasterosteus aculeatus*, which were predominantly detected in the shoreline and shallow transects and not detected in the midline. The winter distribution of these species eDNA was more uniform between transects, with all four species detected in all four transects. Species detection was very similar between the east and west shoreline (which are therefore combined in Figure 3), with the exception that three-spined stickleback were only detected in the east shoreline in summer.

**Figure 3.**
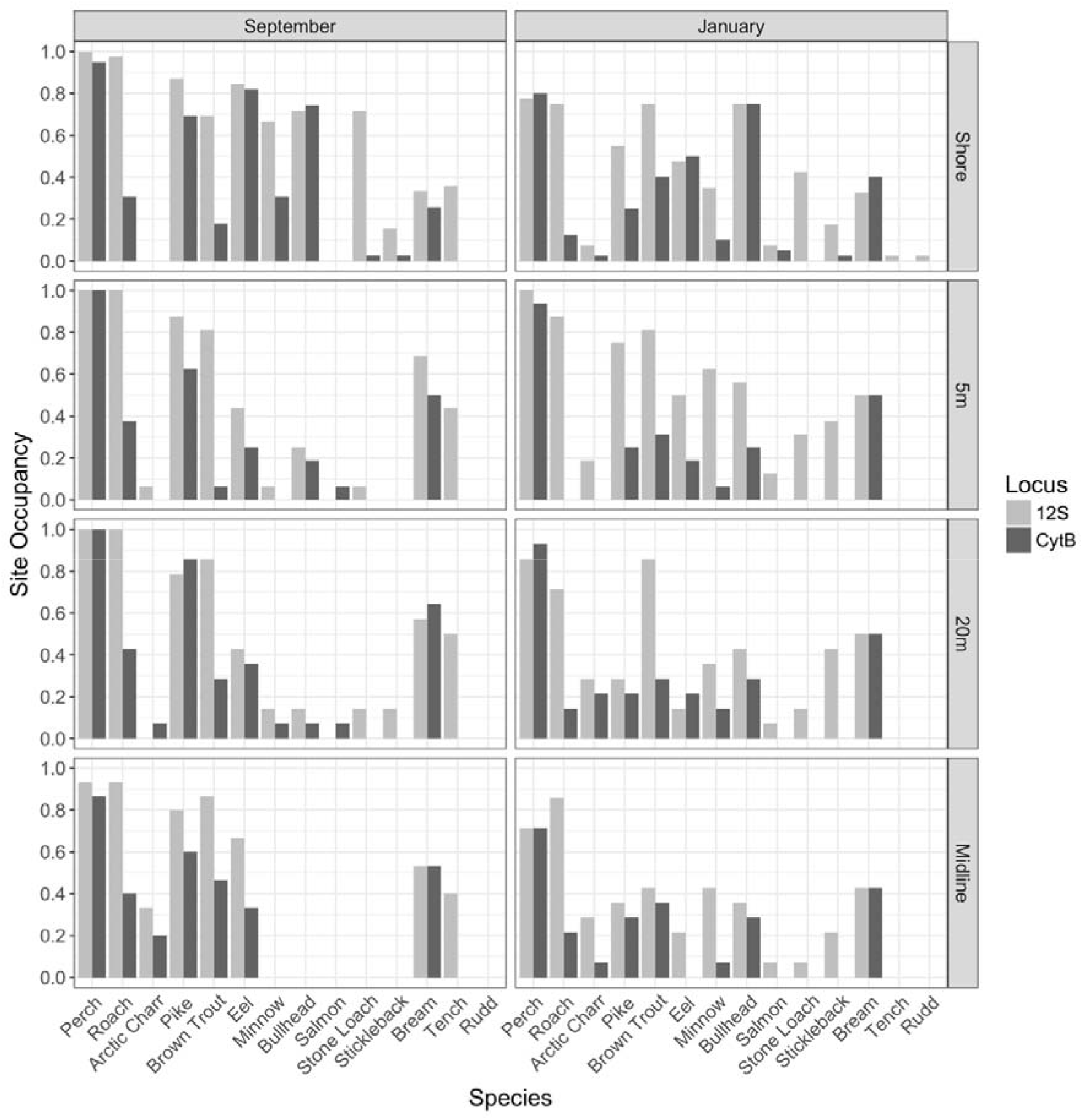
Spatial distribution of eDNA in Windermere for September 2015 and January 2016. Species are ordered according to long term rank. Rows correspond to the four transects: Shore line transect, 5 m transect, 20 m transect and Midline transect (see Figure 1 for details).

A total of 11 species was detected in the ten midline transect sites where depth profiles were taken in September 2015 (Figure 4). The distribution of eDNA at three different depths showed little difference in site occupancy for perch, roach, pike, brown trout, common bream and eel. By contrast bullhead, stickleback and minnow were only detected in the surface water, and Arctic charr was only found in the midwater and bottom sample. Tench was not detected in the epilimnion (but note the detection of tench in other samples may be influenced by contamination as discussed above).

**Figure 4.**
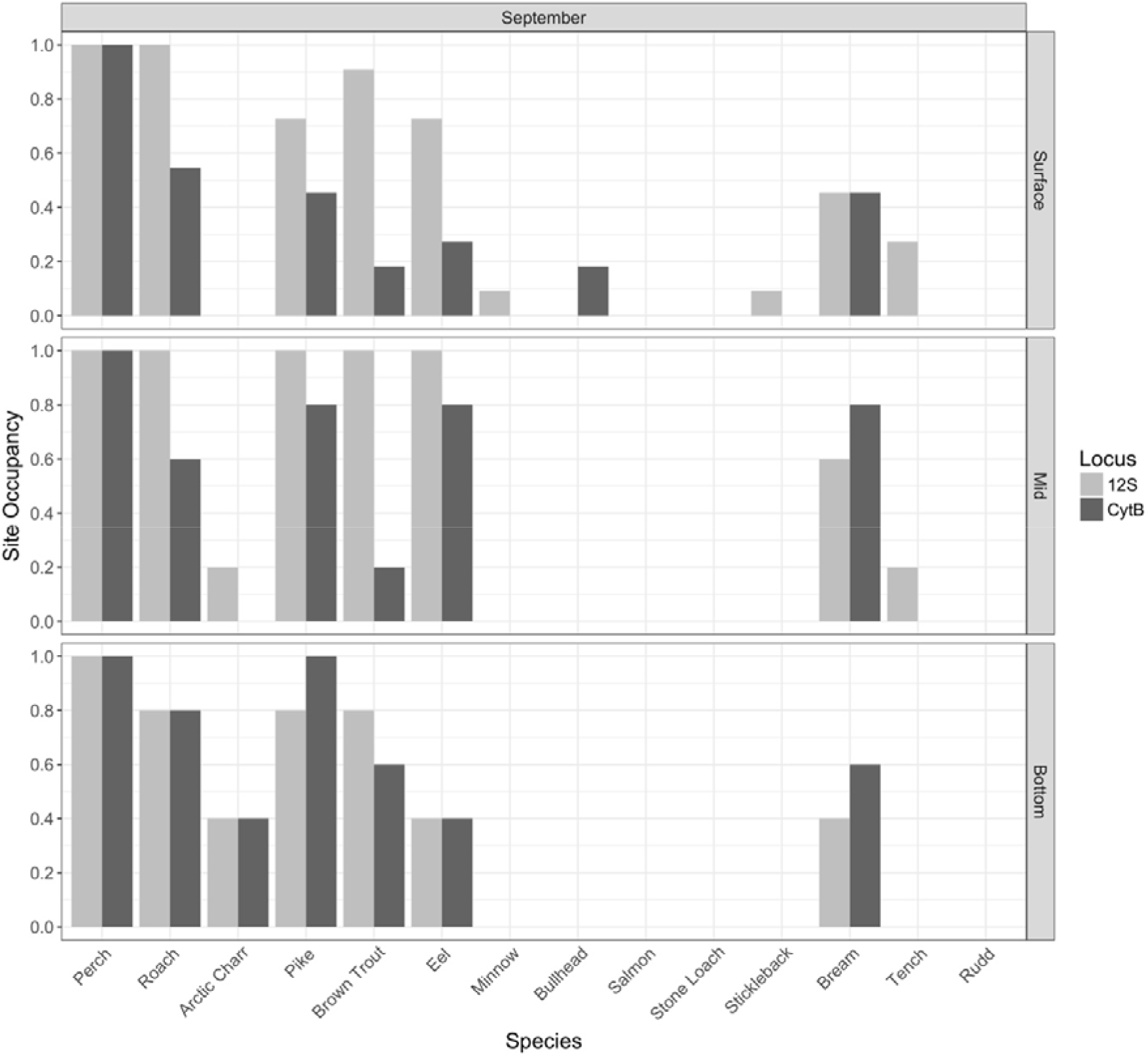
Vertical distribution of eDNA in Windermere from ten sites sampled at the midline in September 2015. Species are ordered according to long term rank. Rows correspond to the three transects: Surface, Mid, and Bottom.

Species accumulation curves based on sample-based rarefaction plateaued consistently higher for 12S than CytB (Figure 5) and curves for shore samples plateaued earlier than for offshore samples in summer (Figure 5a) but not in winter (Figure 5b). The 12S offshore and shore curves plateaued at 10 samples for winter, but in summer around 20 offshore samples were needed to detect the same number of species (Figure 5b). In summer, the 12S shore curve plateaued strongly after 6-10 samples, when 11/14 species (80% of the species diversity) had been captured, whereas the offshore curve continued to increase (Figure 5a). For CytB, offshore and shore curves also start to plateau around 10 samples in winter, when 8-9 species have been captured (57-64% of the species diversity, Figure 5b). In summer, more than 20 shore samples are needed to recover the same number of species detected with just 6 samples sequenced with 12S (Figure 5a).

**Figure 5.**
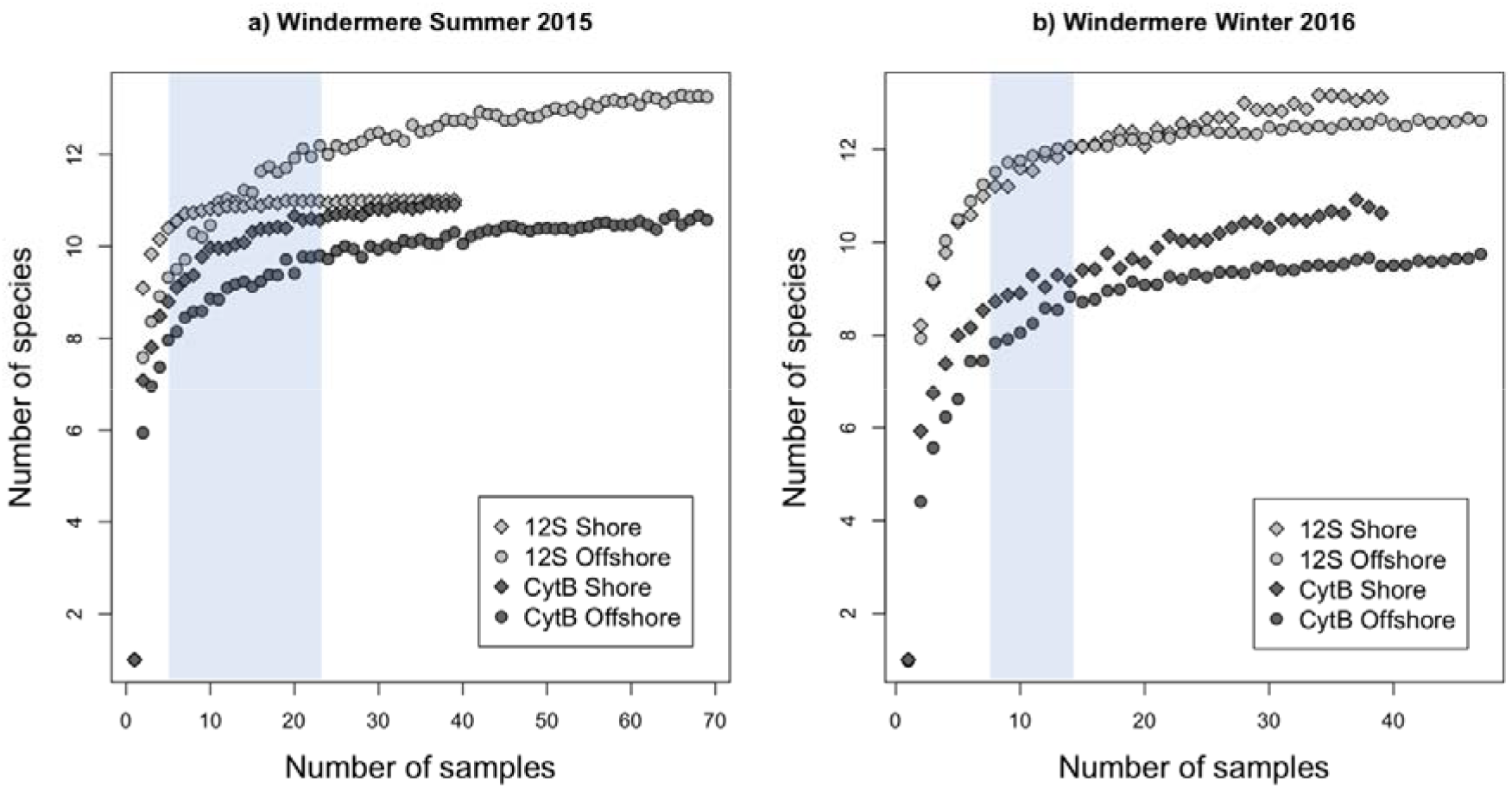
Species accumulation curves based on sample-based rarefaction for Windermere in a) summer 2015 and b) winter 2016. Shore (grey) and offshore (black) samples were analysed separately for 12S (circles) and CytB (diamonds). Shading corresponds to the number of samples needed for optimal species detection.

## Discussion

Few studies have so far explored the spatio-temporal variation in eDNA distribution in aquatic environments. Here, we carried out rigorous spatial sampling in England’s largest lake over three temporal replicates, to determine the level of repeatability in detection and abundance estimation of lake fish species with eDNA metabarcoding. Our analyses demonstrated that species detection and estimation of rank abundance is highly repeatable between seasons, but highlighted some important considerations for design of future fish biodiversity surveys in lakes, which reflect species ecology and seasonal dynamics of aquatic environments.

*eDNA recovers more species than established methods and reflects species relative abundance* In our previous study, carried out in winter 2015, 14 of the 16 species confirmed in Windermere using established methods, were detected using eDNA (Hänfling et al., 2016). The same 14 species were detected in winter 2016, and 13 of the species were detected in September 2015, demonstrating strong consistency in species detection across seasons. By comparison, gill-netting surveys in September 2014 and 2015 found four and five of the most common species respectively (perch, roach, brown trout, pike, in both years, and common bream in 2015). These results add to the growing number of studies that have demonstrated higher detection rates of fish species with eDNA compared to established methods in both freshwater (Valentini et al., 2015; Civade et al., 2016; Hänfling et al., 2016) and marine (Thomsen et al., 2012; Miya et al., 2015; Port et al., 2016; Andruszkiewicz et al., 2017; O’Donnell et al., 2017; Yamamoto et al., 2017) environments.

The only species that were not detected across all sampling campaigns were the river and sea lampreys. We have since detected lamprey eDNA in Windermere and other UK lakes, and can therefore rule out the possibility that our assay is unsuitable. River and sea lamprey are likely to be present in Windermere or its immediate tributaries during September, but they are also likely to be rare and their distributions are probably highly localised due to the very specific lotic habitat requirements of the early life stages of these species (Dawson, Quintella, Almeida, Treble, & Jolley, 2015; Kelly & King, 2001). In a study of sea lamprey distribution in tributaries of the Laurentian Great Lakes, Gingera et al., (2016) found that detection by eDNA was high (81 to 97%) until spawning finished at the end of June, after which it fell to 6% by mid-August. Taken together, these factors could explain their non-detection in the present study. In addition to the lampreys, Rudd, which is the rarest of the 14 species detected with eDNA, and is only present at very low occupancy in South Basin, was not detected during the September sampling. This non-detection could be due to greater spatial structure in the lake during the summer months, as discussed under *“Spatial and seasonal variation in eDNA distribution in Windermere”*; below.

It has recently been argued that sample pooling reduces the detection probability of fish species (Sato, Sogo, Doi, & Yamanaka, 2017), however this is more applicable to studies that pool samples over large spatial scales, and is compensated for in the present study by the high number of samples collected from across the lake. Although the number of false negatives reported here is very low, it might be possible to reduce this even further by increasing the level of replication (Ficetola et al., 2015). In this study we pooled replicates at the sampling (5 × 400 ml volumes) and PCR (3 replicate) stages to reduce the risk of false negatives, while allowing us to sequence a large number of samples within a budget. However sequencing sample replicates separately would allow more accurate estimation of prevalence, detection probability and false positive and negative rates using full site occupancy modelling (Ficetola et al., 2015). This should be considered for future improvements of the method, but there will obviously be a trade off between increasing levels of replication and cost.

Obtaining accurate estimates of abundance from eDNA is thought to be challenging because of the complex dynamics of eDNA in the environment and the large number of opportunities for bias during the experimental work (Barnes et al., 2014; Lawson Handley, 2015). This is particularly true for eDNA metabarcoding (compared to species-specific approaches), in which the number of sequence reads for a particular species can be heavily biased by differential primer binding (primer bias, Deiner et al., 2017; Elbrecht & Leese, 2015) and/or subsampling of species during library preparation (Leray & Knowlton, 2015; Shelton et al., 2016; Deiner et al., 2017). However a growing number of studies have demonstrated significant relationships between abundance estimates generated from eDNA and established data (Hänfling et al., 2016; Thomsen et al., 2016). Building on previous work (Hänfling et al., 2016), we found a consistent, statistically significant relationship between rank abundance (inferred from long-term established data sets) and eDNA data in the form of both site occupancy and normalised read counts. General trends in relative abundance were highly consistent between seasons. The significant relationships demonstrated here are encouraging, but being able to estimate absolute abundance would be preferable to relative abundance. Normalised read counts were positively correlated to biomass of the five species detected in the September 2015 gillnet surveys, but this was driven, at least in part, by brown trout and pike with low biomass and read count, and perch with very high biomass and read count. One possible option to improve estimates of abundance, without relying on correlations, is the addition of internal standard DNAs followed by use of a copy number correction (Ushio et al., 2018). In a recent study of marine fish eDNA, corrected copy numbers were significantly correlated with those obtained by qPCR, providing a promising solution to the low level of confidence in abundance estimation from metabarcoding data (Ushio et al., 2018).

### Spatial and seasonal variation in eDNA distribution in Windermere

Even in lentic water bodies, eDNA is predicted to move away from its source via microcurrents, and this is particularly true in large lakes, which are highly dynamic. Seasonal differences in eDNA distribution are expected in large lakes because of differences in the stratification of the water column between winter and summer. We therefore predicted greater spatial structure - both across the lake surface and at different depths - in eDNA distribution in summer compared to winter.

Firstly, based on our previous results, we predicted a difference in species composition between the North and South Basins of Windermere, which differ in their trophic status. Species that are known to prefer less eutrophic conditions (e.g. Arctic charr, Atlantic salmon, brown trout, minnow and bullhead) were more restricted to the mesotrophic North Basin, while more eutrophic-tolerant species (common bream, roach, rudd, tench and eel) were more common in the eutrophic South Basin (Hänfling et al., 2016). Species that have no clear trophic association were distributed throughout the two basins (stone loach, pike, perch, three-spined stickleback, Hänfling et al., 2016). This demonstrates some spatial structuring even in the winter months, which closely reflects the species ecology. The same broad pattern was confirmed in the summer and winter samples obtained here. In addition, one noteworthy observation is that common bream, which are known to prefer the eutrophic conditions of the South Basin, have lower occupancy in the South Basin in summer relative to winter. The reverse is true for the North Basin, suggesting bream may be migrating into the North Basin during summer months. Whether this pattern is observed on a consistent basis, and if so, determining the underlying ecological triggers warrant further investigation.

Secondly, we predicted a difference in species detection between the shoreline and offshore samples that reflects the species ecology, with greater spatial structuring in the summer months, due to water stratification. eDNA from species that from our earlier study (Hänfling et al., 2016) are expected to be widely distributed in the lake (perch, roach, pike, brown trout, and eel) was detected uniformly between transects in both seasons. However, consistent with our prediction, strong spatial structuring was observed in the summer compared to winter for species that are known to have strict habitat preferences. Most notably, Arctic charr - a deep lake species - was only detected offshore in summer, and at much higher occupancy in the midline compared to shallower 5 m and 20 m transects. This is consistent with Windermere gill-net surveys, which never record Arctic charr inshore outside their late autumn and early winter spawning season. The opposite spatial pattern was found for littoral and benthic species (minnow, bullhead, stone loach and stickleback), which were not detected in the midline transect during the summer and had higher occupancy in the shoreline transect. Three-spined stickleback eDNA was only found along the east shore of the lake in the summer. Distribution of eDNA was far more uniform in the winter samples, with 12 of the 14 species (apart from tench and rudd) detected in all four transects.

Thirdly, we predicted greater spatial heterogeneity in the vertical transects in the summer because of water stratification, compared to winter. A similar result was found to the horizontal transects discussed above in that eDNA for species with an expected wide distribution (e.g. perch, roach, pike, brown trout, common bream and eel) were detected at all three depths, whereas deep-dwelling Arctic charr was only detected in the midwater and bottom samples, and the more littoral and benthic species were only detected in the surface water. This indicates that eDNA is, to some extent, spatially structured within the water column, and that sampling only surface waters during periods of water stratification could miss deep dwelling species. Vertical stratification of eDNA has also been reported in marine environments. For example in a study of species-rich coastal waters of Japan, 50% of 128 coastal marine fish species were detected in both surface and bottom samples, whereas the remaining 50% were detected in either surface or bottom samples (Yamamoto et al., 2017). Similar variation in vertical eDNA distribution has been reported for jellyfish (Minamoto et al., 2017).

Previous studies have demonstrated that eDNA can persist in the environment over relatively large distances (between approximately 2 to 12 km) in natural river systems (Deiner & Altermatt, 2014; Civade et al., 2016), while others have shown eDNA is more patchily distributed in the environment and therefore the likelihood of detecting a target species may decline over short distances between few to hundreds of metres in ponds or small, shallow lakes (Pilliod, Goldberg, Arkle, & Waits, 2013; Eichmiller, Bajer, & Sorensen, 2014) or even coastal environments (Port et al., 2016; O’Donnell et al., 2017; Yamamoto et al., 2017). Our results indicate that the distribution of eDNA within a large, deep lake is patchy, but varies between seasons, with greater heterogeneity in the summer months when lake water is less mixed. Further work is needed to investigate the impacts of microhabitats within the lake and the scale of spatial autocorrelation.

Small, but important differences between seasons and transects, as well as between loci, were demonstrated by the sample-based rarefaction analyses. Previously, based on the winter 2015 sample, we inferred that 10-25 samples detected the majority (≥85%) of the total species detected (Hänfling et al., 2016). The new results broadly support this estimate, but provide important additional insights. Firstly there is a clear difference between the loci in terms of the number of species recovered, with greater power for species detection demonstrated by 12S than CytB. Secondly, species-accumulation curves plateaued earlier for winter than summer, suggesting fewer samples may be needed in winter to detect the same number of species. Approximately 10 samples are needed in winter to recover ≥85% of the species detected, whereas in summer, 10 samples recovers only ~70% of the total species detected by each marker. Finally, there is very little difference between offshore and shore sampling in the winter, in terms of the number of species detected. However there is a notable difference in summer for 12S; the shore curve plateaus strongly at approximately 6 samples, whereas the offshore curve continues to rise. This is consistent with our observations from the transects (i.e. detection of Arctic charr only in deep water during the summer) and indicates that shore sampling only in the summer, may miss species detected at other times of year. To summarise, for our study site, 6-10 shore samples collected in winter and sequenced with 12S are recommended to detect the maximum number of species, with minimum sampling effort.

### Conclusions and recommendations

In summary, we have demonstrated that species detection and estimation of relative abundance of lake fish with eDNA is repeatable between seasons, but there are important spatial and seasonal differences that need to be considered for optimal species detection and abundance estimation. This adds to the growing body of evidence that eDNA is not homogeneously distributed in time or space and can provide an accurate description of aquatic communities (O’Donnell et al., 2017; Macher & Leese, 2017; Stoeckle, Soboleva, & Charlop-Powers, 2017). To maximise the number of species that can be detected, while minimising the costs and effort associated with sampling, we recommend shoreline sampling in the winter and sequencing with 12S, since this assay outperformed CytB in terms of species detection. Following this sampling strategy, 6-10 samples are needed to detect the majority of species known to be present in Windermere. However if abundance estimation is required, it makes more sense to collect as many, spatially representative, samples as possible. Although we found a consistent, significant correlation between rank abundance and eDNA (site occupancy or read count) between seasons, summer sampling, when eDNA is more patchy in distribution, may be preferable (at least in principle) for abundance estimation as site occupancy will more accurately reflect species presence or absence. The minimum number of samples needed to accurately estimate abundance needs to be explored.

## Acknowledgements

This work was funded by the Scottish Environmental Protection Agency (SEPA, contract JUL213921). HJ was funded through the Higher Education Innovation Fund. We are very grateful to Alistair Duguid, Willie Duncan, and Sean Morrison, from SEAP and Kerry Walsh, and Graeme Peirson from the Environment Agency for supporting development of an eDNA based WFD classification tool for lake fish. Ben James and Janice Fletcher helped with the offshore sample collection.

## Data Accessibility

Raw read data for all four libraries have been submitted to NCBI (BioProject: <u>PRJNA482277</u>, SRA Study:<u>SRP154799</u>. https://www.ncbi.nlm.nih.gov/Traces/study/?acc=SRP154799). To assure full reproducibility of our bioinformatics steps, the reference databases and Jupyter notebooks for data processing have been deposited in a dedicated GitHub repository (https://github.com/HullUni-bioinformatics/Handley_et_al_2018).

## Author contributions

LLH, BH, IJW and DSR designed the research and wrote the paper. LLH, BH, DSR, IJW, HK, HJ, JL, CH, RB, RW and RD performed the research. AS helped to develop the bioinformatics pipeline.

